# The effect of anti-IL-6 receptor antibody for the treatment of McH-lpr/lpr-RA1 mice that spontaneously developed ankylosing arthritis

**DOI:** 10.1101/362194

**Authors:** Takuya Izumiyama, Yu Mori, Kazu Takeda, Naoko Mori, Shiro Mori, Tetsuya Kodama, Eiji Itoi

## Abstract

**[Background]:** McH-lpr/lpr-RA1 mice are a new strain of mice which spontaneously develop arthritis in the ankle, leading finally to ankylosis. There is no published data that drug treatment has been trialed on these mice.

**[Objectives]:** This study examined the effect of the mouse anti-IL-6 receptor antibody, MR16-1, for the treatment of ankylosis in McH-lpr/lpr-RA1 mice.

**[Methods]:** Male McH-lpr/lpr-RA1 mice were randomly divided into control and treatment groups. MR16-1 was administered from 10 weeks of age for the treatment group. Saline was applied for the control group. The drug was administered once a week, at an initial dose of 2 mg, then maintained at 0.5 mg once per week thereafter. The effects were evaluated by the histopathological synovitis score, in vivo imaging using indocyanine green liposomes, and analysis of the gene expression of inflammatory cytokines.

**[Results]:** Tissue analyses were carried out at 14, 17 and 20 weeks of age. The synovitis scores of treated groups were significantly lower compared with those of the control group at every age. The kappa coefficient was 0.77. However, progression of ankylosis persisted in the MR16-1 treated group. In vivo imaging using indocyanine green liposomes showed significant decreases in signal intensities of treated groups at week 14, but no significant differences were observed at week 18. Blood serum amyloid A levels in treated groups were significantly lower at 17 weeks of age. The gene expression levels of *Tnf* and *Il17* were also significantly lower in MR16-1 treated groups.

**[Conclusions]:** Administration of the anti-IL-6 receptor antibody is effective for the treatment of synovitis and bone destruction of McH-lpr/lpr-RA1 mice. McH-lpr/lpr-RA1 mice may be a suitable experimental model for the development of new treatments for spondyloarthritis. IL6 signal blockade could contribute to the treatment of spondyloarthritis, and further studies should be carried out to confirm its potential in the prevention of deformity associated with ankylosis.

## Introduction

Spondyloarthritis diseases have complex pathological features, including synovitis, bone erosion, enthesitis and joint ankylosis [1]. The imbalance of bone and cartilage formation and the destruction of joint structure results in the structural changes observed in these diseases. Rheumatoid arthritis is characterized by synovial proliferation and erosion of joint cartilage due to chronic inflammation [2]. On the other hand, spondyloarthritis involves a distinct remodeling process leading to entheseal ossification and joint ankylosis, with synovial proliferation and bone erosion [3,4]. The histological characteristics of ankylosis include the proliferation of cartilage formation, and subsequent replacement of cartilage by bone (endochondral ossification) [5,6]. Recent studies have reported that IL-17 and IL-22 signaling molecules released from IL-23 positive T cells are involved in entheseal chondral proliferation and ossification [7,8]. Patients with ankylosing arthritis are typically treated with tumor necrosis factor (TNF)-α inhibitors. However, the prevention of ankylosis is recognized as being difficult especially in advanced stages of the disease [9-11]. Moreover, animal models of ankylosing arthritis are yet to be established. It is necessary to study animal models of ankylosing arthritis to elucidate the pathological mechanisms of the disease, and develop new treatment methods.

Our research group have previously reported the development of a murine model with spontaneous, progressive ankylosis in ankle joints [12]. Backcross generation mice were prepared using a non-arthritis strain of mice, C3H/HeJ-lpr/lpr (C3H/lpr), MRL/lpr × (MRL/lpr × C3H/lpr) F1. Among these N2 mice, we observed development of arthritis of the ankle joints, with macroscopic swelling. We then began to intercross the N2 mice, by selection based on swelling of the ankle joints. Finally, we established a novel recombinant strain of mice, designated McH-lpr/lpr-RA1, which showed a high incidence of arthritis with enthesopathy. Analysis of the joint remodeling processes may elucidate the fundamental mechanisms of spondyloarthritis and other ankylosing diseases.

The anti-IL-6 receptor monoclonal antibody is available for clinical use in patients with rheumatoid arthritis, and has been shown to have therapeutic effects in the treatment of severe rheumatoid arthritis [13]. The effect of anti-IL-6 treatment in patients with ankylosing spondylitis has also been reported — studies revealed that the anti-IL-6 receptor antibody did not prevent the progression of spinal changes associated with ankylosis [14]. Furthermore, the effect of treatment with anti-IL-6 receptor antibody on peripheral symptoms of spondyloarthritis has also been reported, but its use in joint ankylosis remains unclear [15]. MR16-1 is a rat anti-IL-6 receptor monoclonal antibody, as previously described in the literature [16]. MR16-1 is used in many experimental disease models to assess the effects of blocking IL-6 signaling [17-21]. McH-lpr/lpr-RA1 mice may provide a suitable animal model to assess the therapeutic effects of IL-6 signal blockade in ankylosing arthritis, and the administration of MR16-1 will determine the potential treatment effects of IL-6 ligand and receptor blockade.

The purpose of the present study is to investigate the effect of MR16-1 for the treatment of ankylosing arthritis, and the mechanism of inflammatory change and ankylosis in the experimental murine model of McH-lpr/lpr-RA1 mice. We hypothesized that IL-6 signal blockade would suppress the progression of synovial proliferation and joint ankylosis in McH-lpr/lpr-RA1 mice.

## Methods

### Animals

McH/lpr-RA1 mice were generated using F54 C3H/HeJ-*lpr*/*lpr* and MRL/lpr × (MRL/lpr × C3H/lpr) mice. This recombinant congenic strain of mice was designated McH/lpr-RA1 as previously described in the literature [12]. All mice were housed in the animal unit of Tohoku University Medical School, an environmentally controlled and specific pathogen-free facility. Animal protocols were reviewed and approved by the Tohoku University Animal Studies Committee. All experiments were performed using week 10 male mice.

### Treatment of mice

IL-6 signal blockade was performed with an intraperitoneal injection of 2 mg of rat anti-mouse IL-6R mAb (MR16-1, a kind gift from Chugai Pharmaceutical, Tokyo, Japan), once in the first treatment (week 10). Thereafter, 0.5 mg of MR 16-1 was administered once a week until 20 weeks of age as previously described in the literature [22] Phosphate buffered saline (PBS) was administered on the same schedule as a negative control.

### Microcomputed tomography analysis

Microcomputed tomography (micro-CT) imaging was performed at 20 weeks of age (n=5 for each group). Harvested tibiae were stored in 70% ethanol at 4 °C, and analyzed using a micro-CT scanner (Scan Xmate-L090; Comscan Techno Co. Ltd., Kanagawa, Japan) operated at a peak voltage of 75 kV and 100 μA. The scanned region included 505 images from the proximal end of the tibia, at a resolution of 10.4 μm per voxel, and an image size of 516 × 506 pixels. Bone volume (BV; mm^2^), total volume (TV; mm^2^), bone volume fraction (BV/TV; %) and trabecular thickness (mm) were evaluated and calculated from the axial slice of the proximal tibia using TRI/3D-BON software (Ratoc System Engineering Co., Tokyo, Japan), as previously described in the literature [23].

### Enzyme-linked immunosorbent assay

Serum amyloid A (SAA) and IL-6 levels were determined using an enzyme-linked immunosorbent assay (ELISA) kit for SAA and IL-6 (Biosource, Camarillo, CA and R&D Systems Inc., Minneapolis, MN, USA) according to the manufacturer’s recommendations at 14 and 17 weeks of age (n=5 for each group). Briefly, serum samples were diluted 1:200 in assay diluent and incubated with conjugated anti-mouse SAA antibody. Serum samples were incubated with anti-mouse IL-6 antibody without dilution. Substrate tetramethylbenzidine was added, samples were read at OD450 nm and results were analyzed using the four-parameter fit to determine SAA values.

### Histomorphometric analysis

Ankle joints were harvested at 14, 17 and 20 weeks of age and decalcified by soaking in 220 mM/L EDTA-Na for 3 weeks. Decalcified samples were sectioned and stained with hematoxylin and eosin for histopathological evaluation of synovitis and ankylosis, as previously described in the literature [24]. The calculated synovitis score was the sum of scores for: enlargement of the synovial lining cell layer (0 points: thickness of 1 layer; 1 point: thickness of 2-3 layers; 2 points: thickness of 4-5 layers; 3 points: thickness of more than 5 layers), density of the resident cells (0 points: normal cellularity; 1 point: slightly increased cellularity; 2 points: moderately increased cellularity; 3 points: greatly increased cellularity, pannus formation and rheumatoid like granulomas might occur) and inflammatory infiltrate (0 points: no inflammatory infiltrate; 1 point: few lymphocytes or plasma cells; 2 points: numerous lymphocytes or plasma cells, sometimes forming follicle-like aggregates; 3 points: dense, band-like inflammatory infiltrate, or numerous large, follicle-like aggregates). The synovitis score was assessed by two researchers and the kappa coefficient was calculated (n=5 for each group).

### Synthesis of indocyanine green liposomes

Indocyanine green (ICG) liposomes were prepared as previously described in the literature [25-27]. Briefly, 1,2-distearoyl-sn-glycero-3-phosphatidylcholine (DSPC; NOF, Tokyo, Japan) and 1,2-distearoyl-sn-glycero-3-phosphoethanolamine-N-[methoxy-(polyethylene glycol)-2000] (DSPE-PEG[2000-OMe]) (DSPE-PEG:NOF, 94:6 mol/mol) were placed in a pear-shaped flask, and chloroform was added until the lipids were completely dissolved. The chloroform was then evaporated under reduced pressure using a rotary evaporator (NVC-2100/N-1000, Eyela, Tokyo, Japan) until a lipid film remained.

Next, 100 μM of ICG (Daiichi Sankyo, Tokyo, Japan) dissolved in 10 mL PBS was added to the thin lipid film to form multi-lamellar liposome vesicles. After repeated freeze-thaw cycles, the size of the liposomes was adjusted to <100 nm using extrusion equipment (Northern Lipids Inc., Vancouver, BC, Canada) with four filter sizes (100, 200, 400 and 600 nm; Nuclepore Track-Etch Membrane, Whatman plc, Maidstone, UK). For sterilization, liposomes were passed through a 0.45 μm pore size filter (Millex HV filter unit, Durapore polyvinylidene-difluoride [PVDF] membrane, EMD Millipore, Billerica, MA, USA). Unbound ICG was removed using PD-10 columns. The lipid concentration was measured using the Wako Phospholipids C test (Wako Pure Chemical Industries, Osaka, Japan), and adjusted to 1.0 μmol/mL. Diameters of the ICG liposomes were measured using a particle size analyzer (ELSZ-2,Otsuka Electronics, Osaka, Japan).

### In vivo bioluminescence imaging system analysis

In vivo bioluminescence imaging system (IVIS) analysis was performed at 10, 14 and 18 weeks of age in MR16-1 treated and control mice (n=5 for each group). Two hundred milliliters of ICG liposomes was administered intravenously. The fluorescence intensity of leaked ICG was measured at 0, 5, 30, 60, 120 and 180 minutes after injection using an IVIS; Xenogen, Alameda, CA, USA), as previously described in the literature [25,26].

### Quantitative RT-PCR

Synovial tissue specimens from ankle joints were harvested at 10, 14 and 16 weeks of age (n=5 for each group). Total RNA was extracted using QIAzol Lysis Reagent (Qiagen, Hilden, Germany) and an RNeasy Mini Kit (Qiagen, Hilden, Germany). Synthesis of cDNA from total RNA was carried out using RT buffer, RT random primers, dNTP mix, and Multiscribe reverse transcriptase (Applied Biosystems, Foster city, CA, USA). A total of 9 μL cDNA diluted 1:9 was added to 10 μL Taqman Universal Master Mix II with Uracil N-glycosylase (Applied Biosystems, Foster City, CA, USA). Real-time amplification of the genes was performed using 1 μL ready-to-use Taqman Gene Expression Assays (Applied Biosystems) for *Il6*, *Tnf*, *IL17,* and *gapdh* as an endogenous control (assay IDs: Mm00446190_m1, Mm00443260_g1, Mm00439618_m1, and Mm99999915_g1). Relative gene expression data were analyzed using the delta-delta-Ct method with PCR-efficiency correction using StepOne software version 2.2.2 (Applied Biosystems), as previously described in the literature [28].

### Statistical analysis

Statistical analyses were performed using JMP software version 13.1 (SAS, Cary, NC, USA). All data are expressed as the mean ± standard error (SE). Statistical significance of the differences between values were evaluated using the Student’s t-test. Values of p <0.05 were regarded as statistically significant. Kappa coefficients were calculated using SPSS version 21 (IBM, Armonk, NY, USA).

## Results

### Micro CT analysis

In the coronal-reconstructed micro CT images at 20 weeks of age, there were no apparent differences between MR16-1 treatment group and control group mice (Fig 1A). Quantitative structural analyses of proximal tibia are shown in Fig 1B. No significant differences were seen in BV, TV, BV/TV or trabecular thickness between the two groups. MR16-1 treatment therefore does not correlate with an increase in bone volume in McH/lpr-RA1 mice.

**Figure 1.**
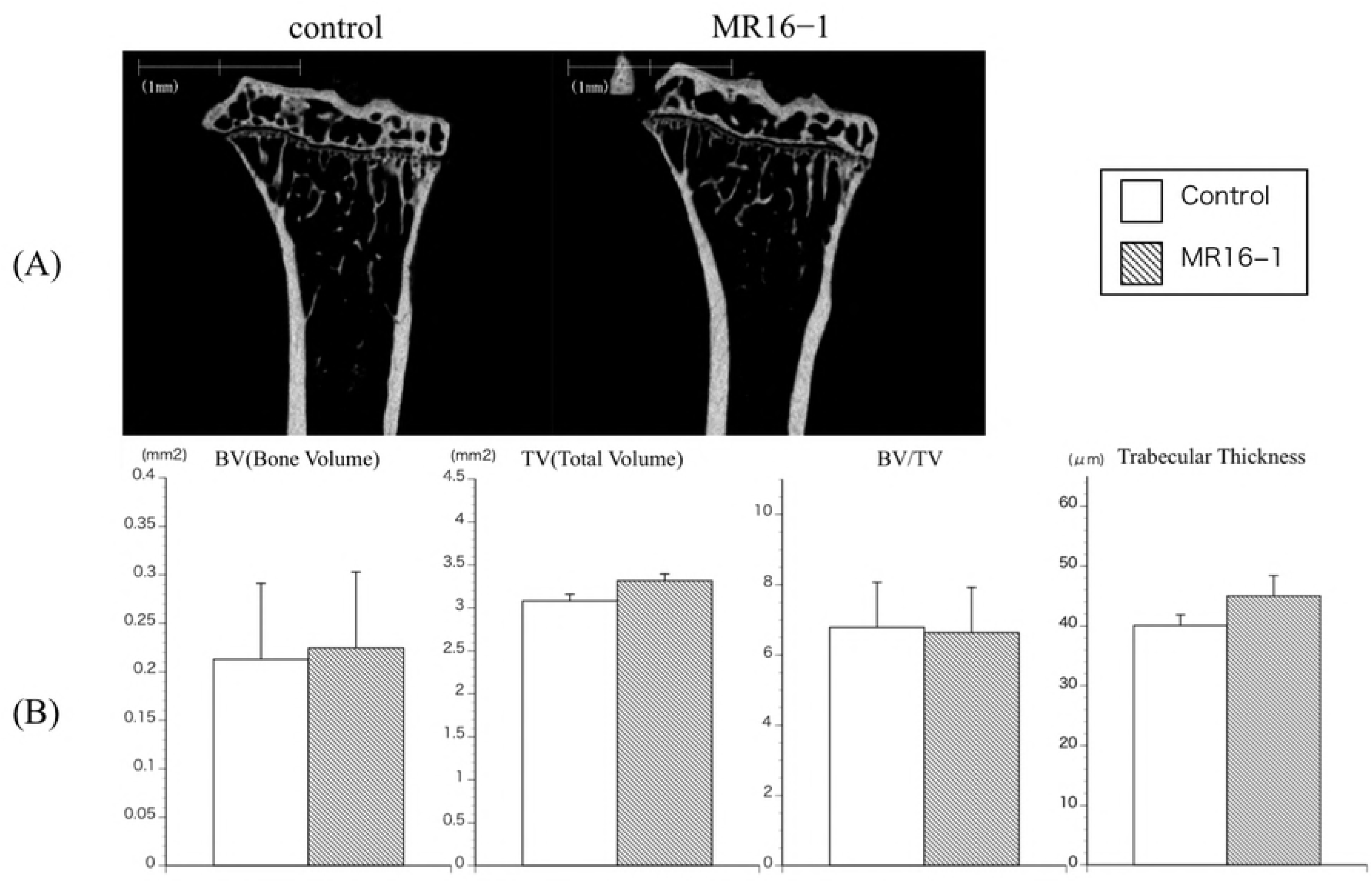
Micro CT image of tibia at week 20 in control and MR16-1 treated groups. A. Representative coronal reconstruction images of proximal tibia in control and MR16-1 treated groups.
B. Structural parameter analyses of tibia of control and MR16-1 treated mice. There are no significant differences in BV, TV, BV/TV and trabecular thickness.

### ELISA assay

The results of the ELISA assays showed reduced levels of SAA in the MR16-1 treatment group at 14 weeks of age, with significantly decreased levels at week 17 (p =0.0011) (Fig 2A). The results of the assay for serum IL-6 indicated no significant difference between the control and MR16-1 treatment groups at 14 and 17 weeks of age (Fig 2B).

**Figure 2.**
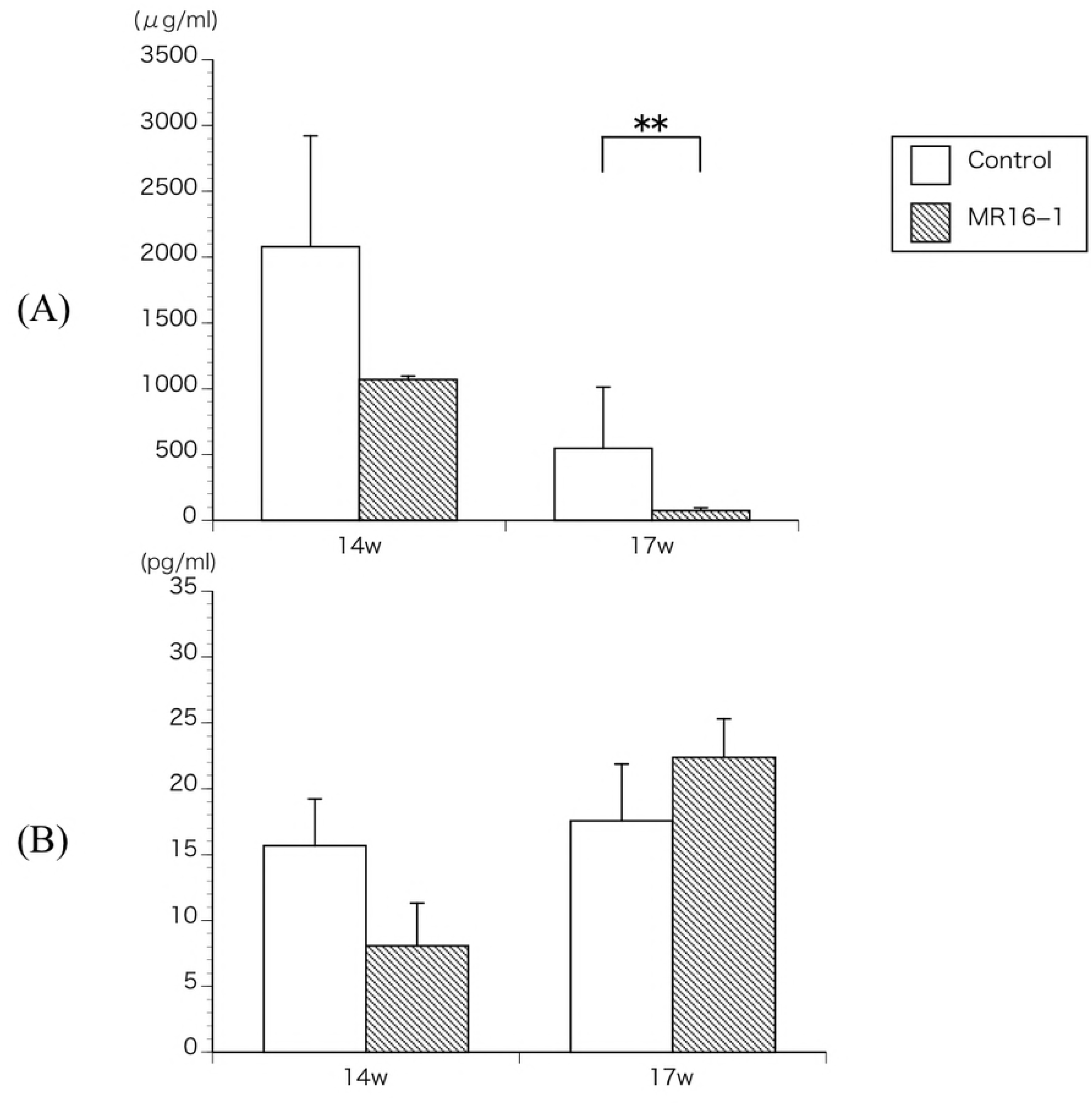
ELISA assays for SAA and IL-6. A. ELISA assay for SAA at week 14 and 17. Serum SAA level is significantly lower in MR16-1 treated group at week 17.
B. ELISA assay for IL-6 at week 14 and 17. There is no significant difference in serum IL-6 level.

### Histomorphometric analysis

At 14 weeks of age, control group mice showed apparent synovitis, pannus formation, entheseal inflammation and ankylosis in the ankle region (Fig 3A and 3B). In MR16-1 treated mice, synovitis and pannus formation was suppressed compared with the control group (Fig 3C and 3D). At 17 weeks of age, control group mice exhibited acceleration of synovitis, pannus formation and ankle joint ankylosis (Fig 4A and 4B). In the MR16-1 treatment group, no synovitis or pannus formation were detected, however, the progression of entheseal ankylosis was seen (Fig 4C and 4D). At 20 weeks of age, control group mice showed degenerative synovial lesions, advanced pannus formation and ankylosing change (Fig 5A and 5B). In the MR16-1 treatment group, ossification and ankylosis of the entheseal area was found, although synovitis and pannus formation were absent (Fig 5C and 5D). Histopathological assessments of ankle arthritis were performed by calculation of the synovitis score. The Kappa coefficient was calculated to be 0.77, and the reproducibility was good. At week 14, the synovitis score of the control group was 4.29 ± 019, and that of the MR16-1 group was 2.75 ± 0.53. This indicates a significant difference between the two groups (p = 0.0036). At week 17, the synovitis score of the control group was 3.95 ± 0.22, and that of the MR16-1 group was 1.64 ± 0.27. This indicates a significant difference between the two groups (p <0.001). At week 20, the synovitis score of the control group was 3.33 ± 0.28, and that of the MR16-1 group was 2.25 ± 0.3. This indicates a significant difference between the two groups (p =0.0125) (Fig 6).

**Figure 3.**
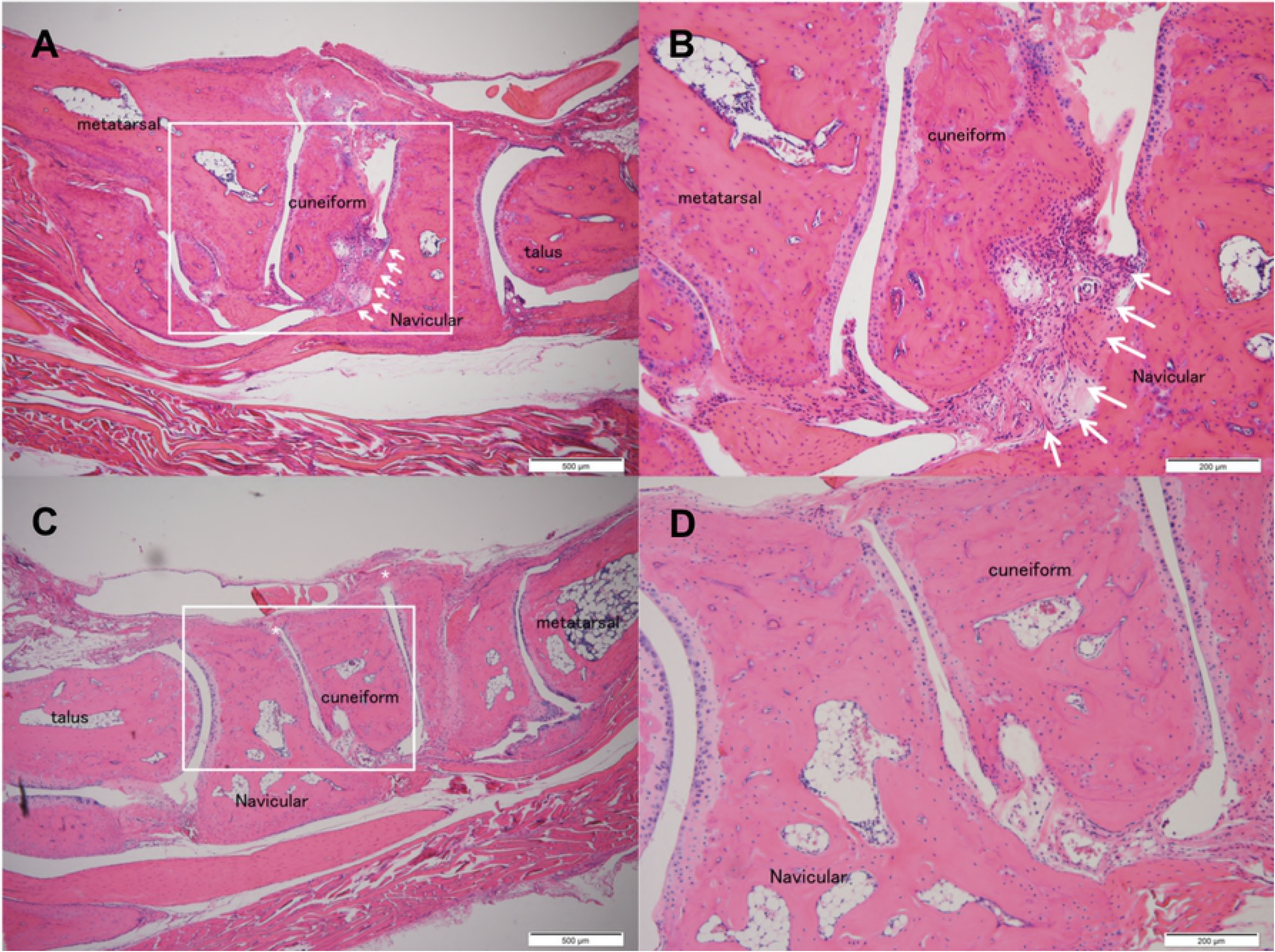
Histological images of control and MR16-1 treated groups at week 14. Representative histological images of ankle joint and foot in control group. (A) is ×40 lower magnification image. (B) is ×100 higher magnification image. Rectangle shows the higher magnification area. Arrows indicate synovial proliferation. ^∗^ indicates entheseal fibrous and cartilaginous ankylosis. Representative histological images of ankle joint and foot in MR16-1 treatment group. (C) is ×40 lower magnification image. (D) is ×100 higher magnification image. Rectangle shows the higher magnification area. Arrows indicate synovial proliferation. ^∗^ indicates fibrous and cartilaginous ankylosis.

**Figure 4.**
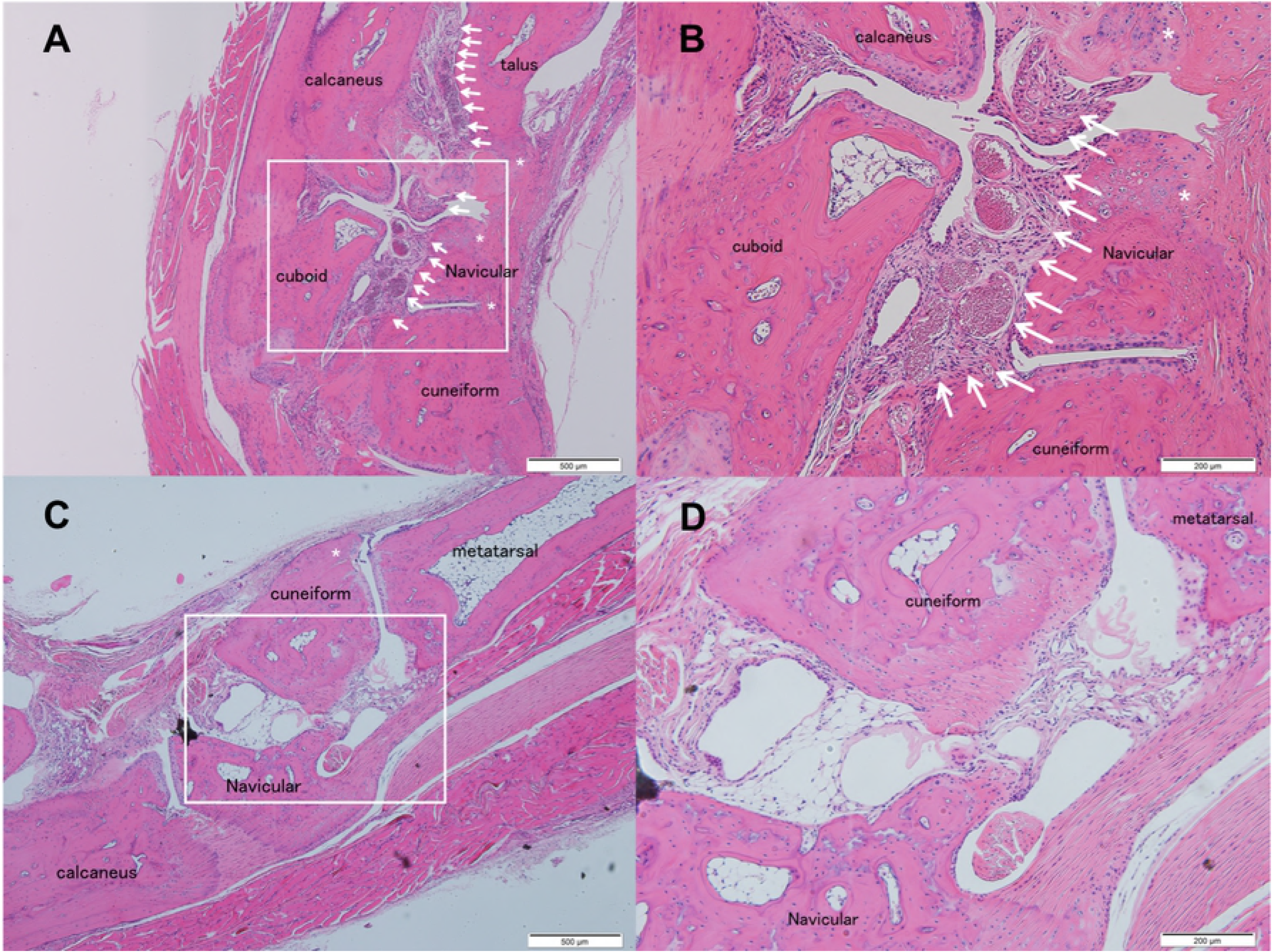
Histological images of control and MR16-1 treated groups at week 17. Representative histological images of ankle joint and foot in control group. (A) is ×40 lower magnification image. (B) is ×100 higher magnification image. Rectangle shows the higher magnification area. Arrows indicate synovial proliferation. ^∗^ indicates entheseal fibrous and cartilaginous ankylosis. Representative histological images of ankle joint and foot in MR16-1 treatment group. (C) is ×40 lower magnification image. (D) is ×100 higher magnification image. Rectangle shows the higher magnification area. Arrows indicate synovial proliferation. ^∗^ indicates entheseal fibrous and cartilaginous ankylosis.

**Figure 5.**
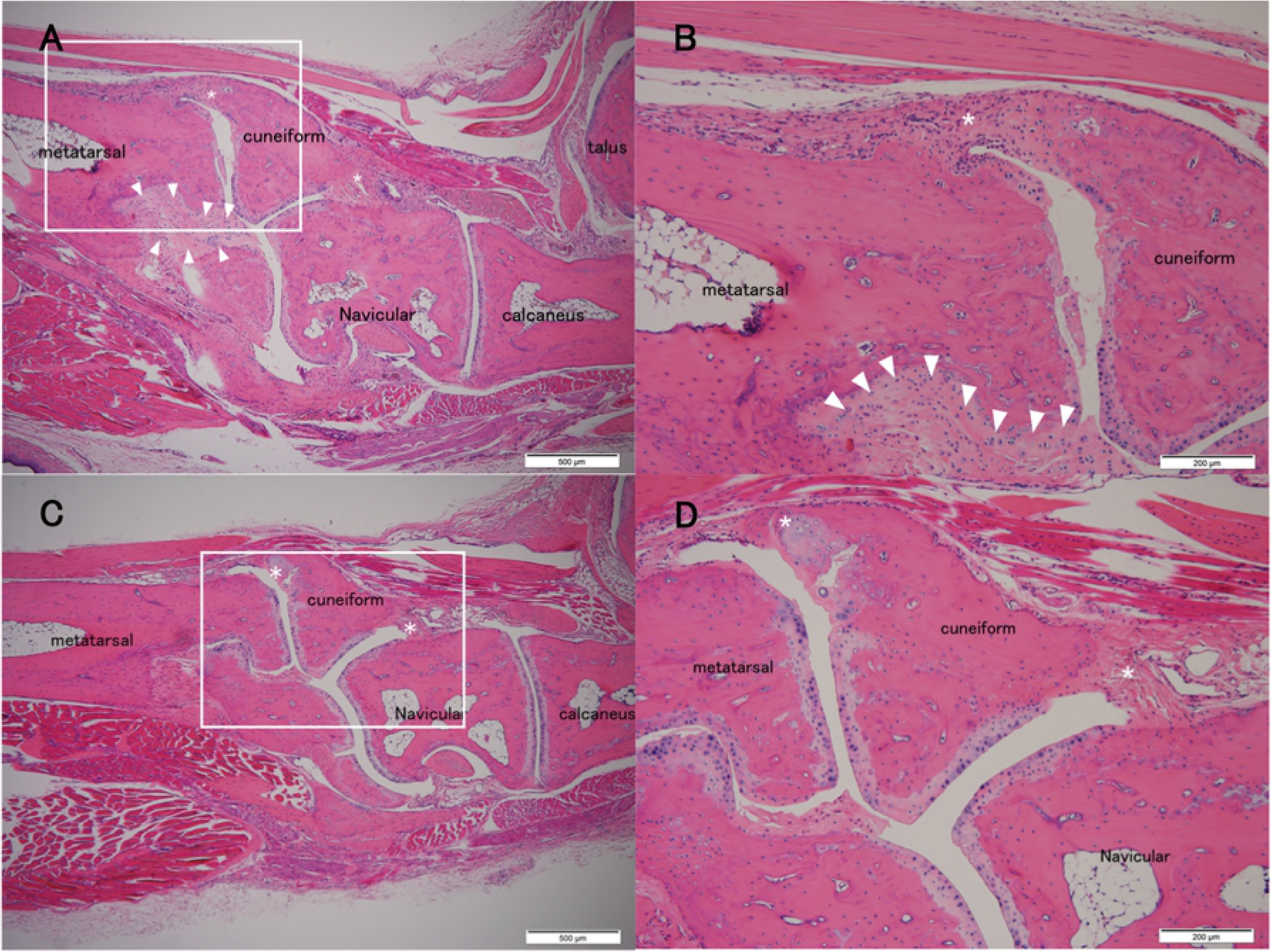
Histological images of control and MR16-1 treated groups at week 20. Representative histological images of ankle joint and foot in control group. (A) is ×40 lower magnification image. (B) is ×100 higher magnification image. Rectangle shows the higher magnification area. Arrow heads indicate subsequent interosseous ankylosis after synovial proliferation. ^∗^ indicates entheseal fibrous and cartilaginous ankylosis. Representative histological images of ankle joint and foot in MR16-1 treatment group. (C) is ×40 lower magnification image. (D) is ×100 higher magnification image. Rectangle shows the higher magnification area. ^∗^ indicates entheseal fibrous and cartilaginous ankylosis.

**Figure 6.**
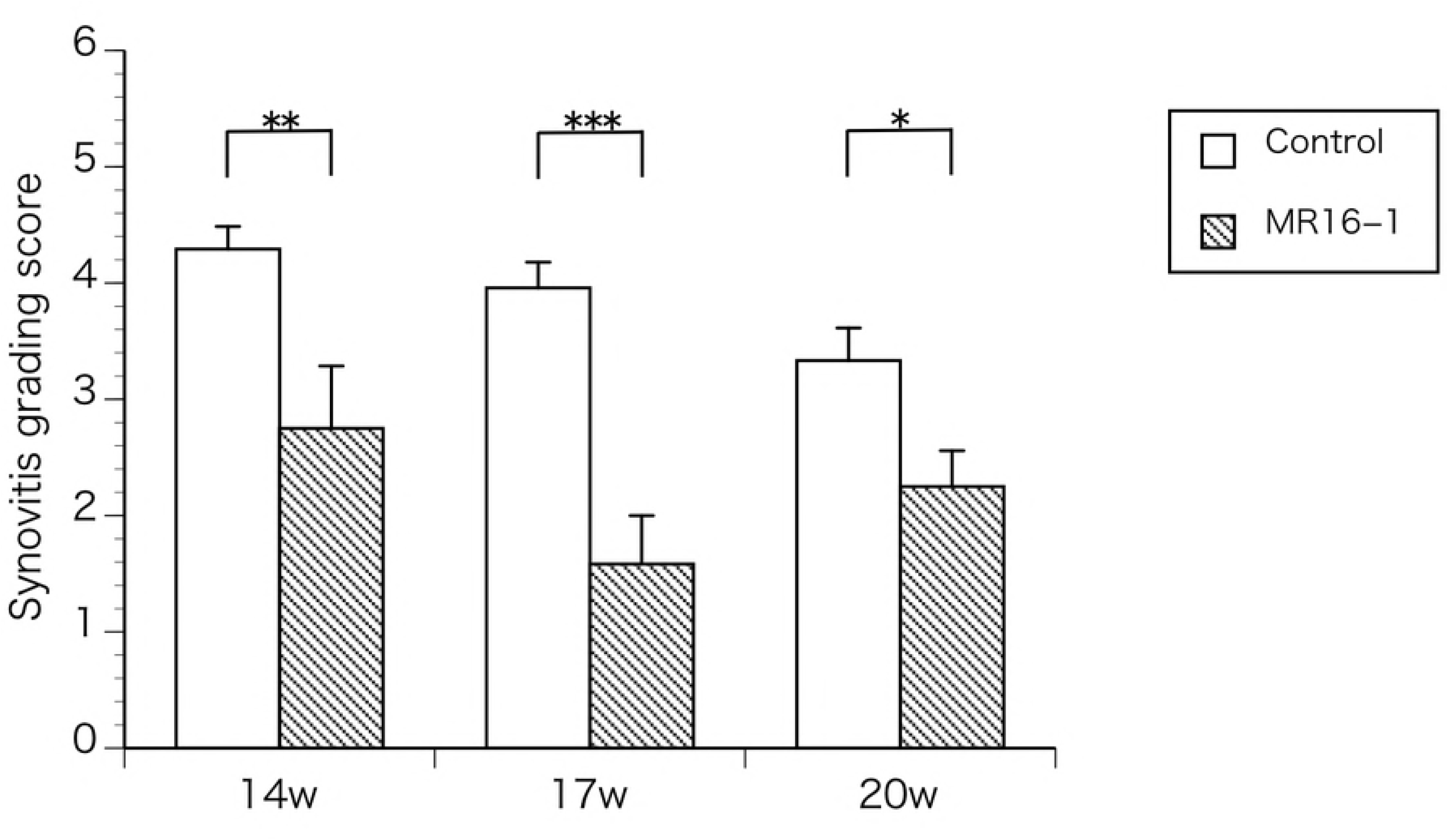
Histomorphometric analyses with the synovitis grading score of foot and ankle joints in control and MR16-1 treated mice. The synovitis grading scores of MR16-1 treatment group are significantly lower in week 14, 17 and 20 compared with those of control group. ^∗^ < 0.05, ^∗∗^ < 0.01, ^∗∗∗^ < 0.001.

### IVIS analysis

The changes in the signal intensities pre-injection and 120 minutes after injection were assessed. Representative images of 14 weeks of age control mice and MR16-1 treated mice are shown in Fig 7A and 7B. MR16-1 treated mice did not show ICG accumulation in the ankle or foot joints. However, control mice showed obvious accumulation in the foot and ankle joints. The results of IVIS analyses are shown in Fig 8. Quantitative IVIS was carried out at weeks 10, 14 and 18 in control and MR16-1 treated mice. There was a significant difference in the signal intensities of control and MR16-1 treated mice at week 14 (p=0.0003), although there no significant differences were seen at weeks 10 or 18.

**Figure 7.**
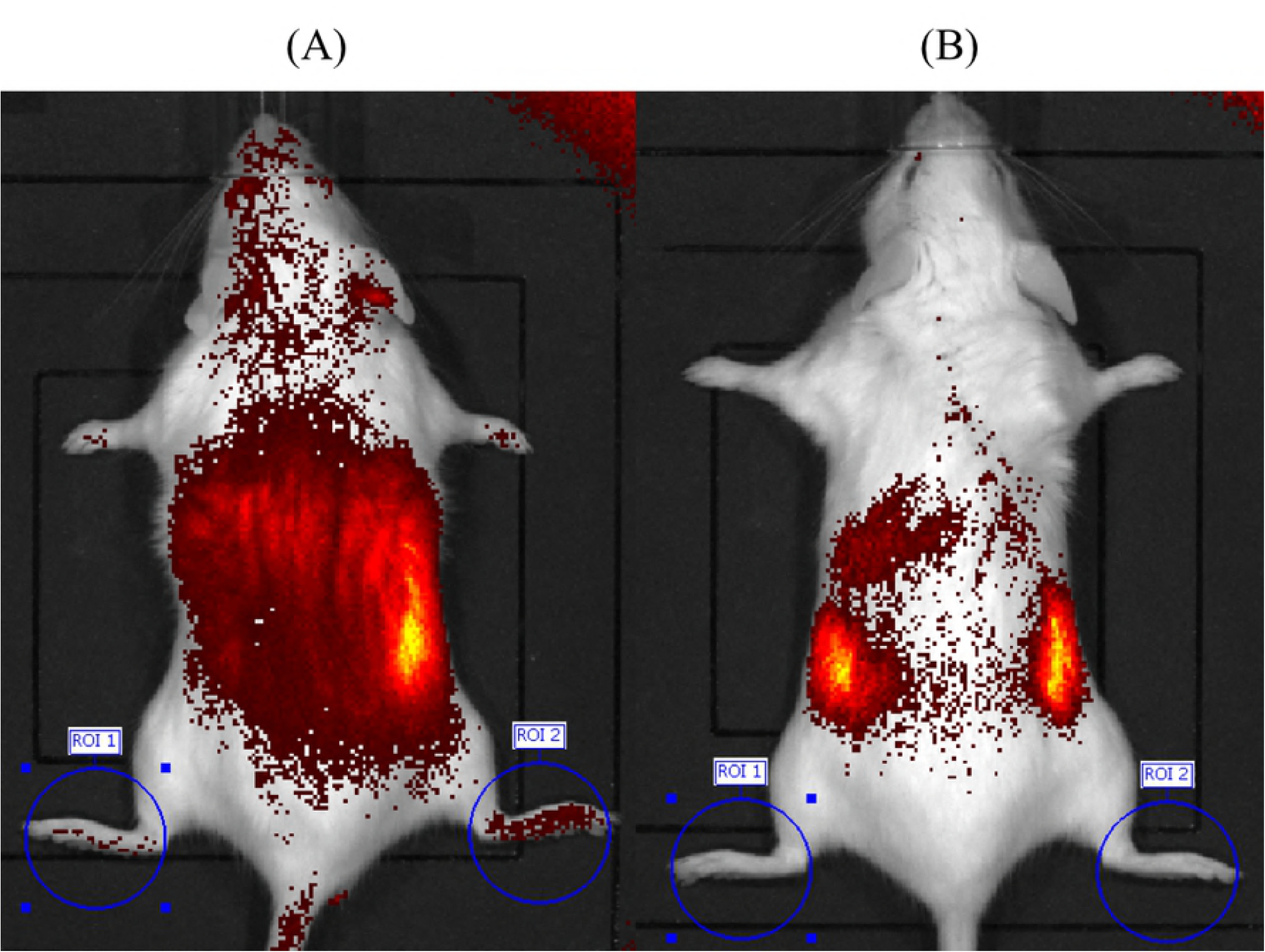
IVIS analyses in control and MR16-1 treated mice. A. Representative image of IVIS of control mouse at week 14. There are evident signals in both ankles and foots.
B. Representative image of IVIS of MR16-1 treated mouse at week 14. There is no significant uptake signals in ankles and foots.

**Figure 8.**
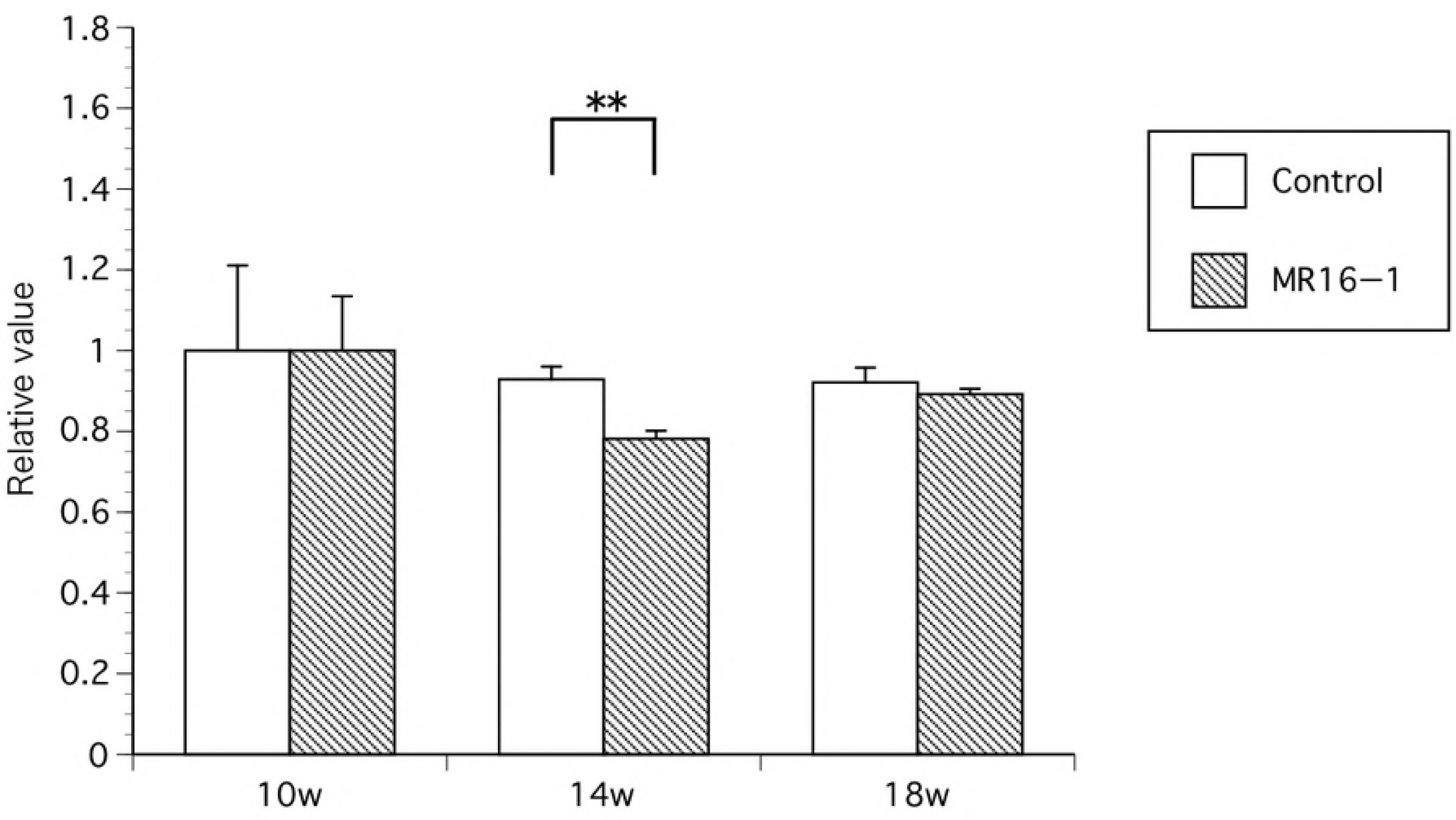
Quantitative analyses of signal intensity of IVIS at week 10, 14, 18. At week 14, the signal intensity of IVIS of ankles and foots is significantly lower in MR16-1 treatment group, compared with the control group. There are no significant changes in week 10 and 18 in both groups. ^∗∗^ p < 0.01

### Quantitative RT-PCR

The results of quantitative PCR analyses are shown in Fig 9. *Tnf* gene expression gradually decreased until week 16 in the MR16-1 treatment group. There was a significant difference in *Tnf* gene expression between the two groups at week 16 (p=0.004), which was lower in the MR16-1 treated group than the control group. *Il17* gene expression also gradually decreased until week 16 in the MR16-1 treatment group. There were also significant differences in *Il17* gene expression between the two groups at week 14 and 16 (p=0.044; p=0.042), which were lower in MR16-1 treated group. *Il6* gene expression was also evaluated, revealing no significant difference in expression levels between the groups.

**Figure 9.**
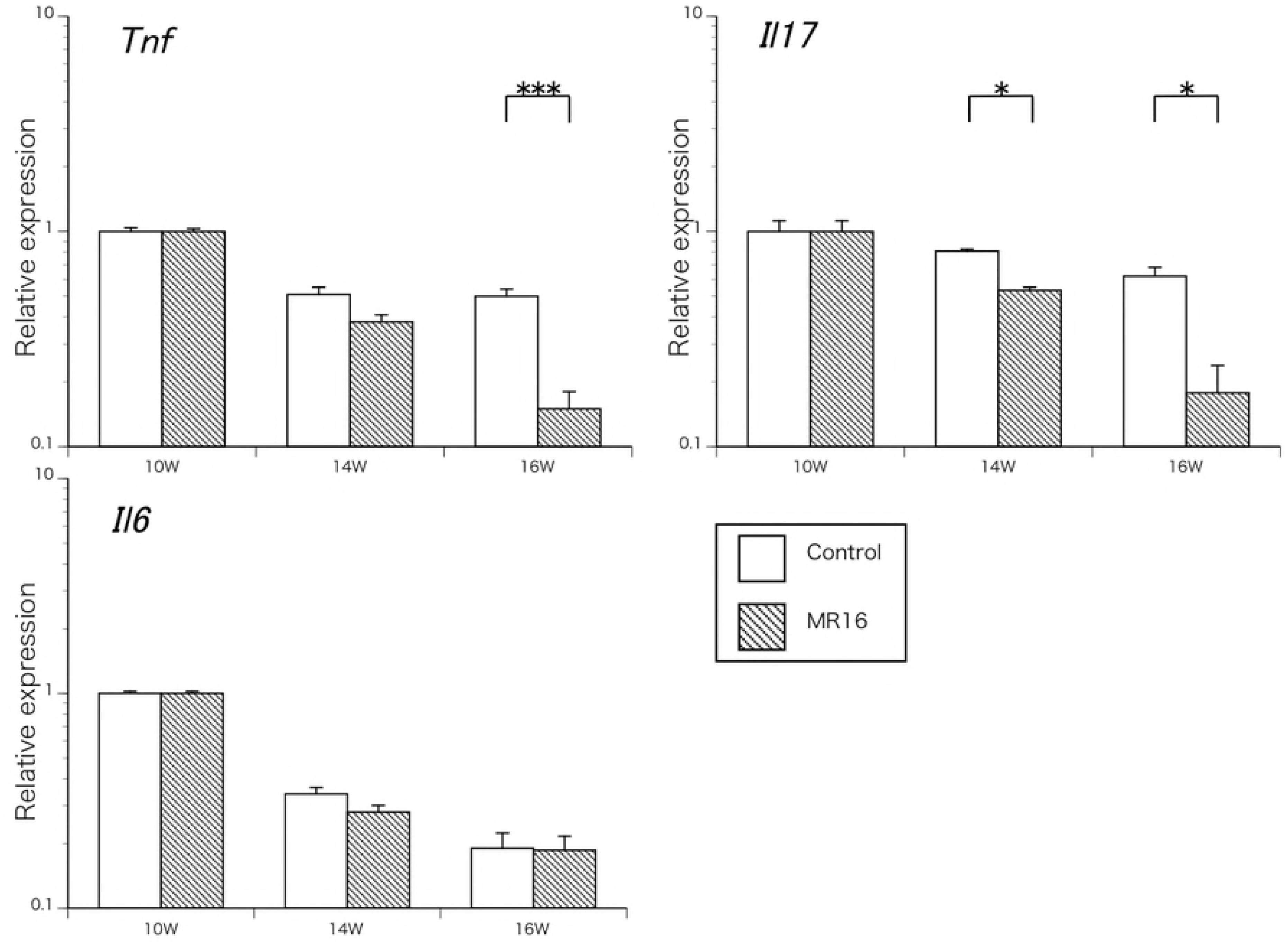
mRNA expression of *Tnf*, *Il17* and *Il6* genes. The expression levels of *Tnf* is significantly lower in MR16-1 treated group at week 16. The expression levels of *Il17* is significantly lower in MR16-1 treated group at week 14 and 16. There is no significant difference of *Il6* expression among both groups. ^∗^p < 0.05, ^∗∗∗^p < 0.001

## Discussion

Previous studies have reported several murine models that develop spontaneous ankylosis [29]. For example, DBA/1 mice spontaneously develop ankylosing arthropathy in the ankle joints [30-32]. However, these mice mainly exhibit fibrous proliferation in enthesis and fibrous ankylosis, without erosion and bone destruction [29]. McH/lpr-RA1 mice show synovitis, pannus formation and ankylosis, resembling spondyloarthritis [12]. We consider that McH/lpr-RA1 mice are a suitable experimental disease model of spondyloarthritis. The present study explores the effects of treatment with the anti-IL-6 receptor monoclonal antibody in McH/lpr-RA1 mice. In the MR16-1 treated group, results of histomorphometric and IVIS analyses showed that the proliferation of synovium and joint destruction were suppressed. However, histological investigations confirmed that ankylosis progressed even with IL-6 blockade treatment. We propose that the anti-IL-6 receptor antibody contributes to the suppression of synovial proliferation in McH/lpr-RA1 mice, but that further factors are involved in the prevention of entheseal ankylosis.

Changes in the bone parameters of MR16-1 treated and control mice were evaluated by micro-CT imaging. In this study there were no significant differences in the bone parameters between two groups. A published study reported the treatment effects of MR16-1 to include increased bone volume in DBA/1 mice [33]. However, in this study McH/lpr-RA1 mice did not show increased bone volume following MR16-1 treatment. In the published study, the dose of MR16-1 was much higher — 8 mg was administered to DBA/1 mice. Our different findings of the effect of MR16-1 treatment on bone volume were likely due to the different dosage of MR16-1.

SAA was significantly lower in the MR16-1 treated group, and we consider that the dose of MR16-1 that was administered is sufficient to suppress inflammation in McH/lpr-RA1 mice. MR16-1 is an anti-IL-6 receptor antibody for the treatment, so it is unsurprising that there were no changes in serum IL-6 level. The results of quantitative RT-PCR showed that gene expression levels of *tnf* and *IL17* were suppressed by IL-6 signal blockade. Previous studies reported that IL-6 signal blockade by MR16-1 suppresses IL-17 signaling [34,35]. Suppression of IL-6, TNF-α and IL-17 signals may contribute to the prevention of synovitis and bone destruction in McH/lpr-RA1 mice. However, only partial prevention of joint ankylosis was seen in the MR16-1 treated group — at week 20 the histological images showed the progression of ankylosis, regardless of the prevention of synovial proliferation and bone destruction. In the results of previous clinical trials, administration of anti-IL-6 receptor antibody was insufficient for complete treatment of ankylosing spondylitis [14]. Some studies have reported that entheseal ankylosis in spondyloarthritis is related to IL-17 signaling [36]. The blockade of IL-17 signaling was achieved in this study; however, this signal suppression might be insufficient, as other signaling pathways (such as IL-12, IL-22 and IL-23) could be involved in the mechanisms of entheseal ossification and ankylosis. Our results are consistent with the results of human clinical trials of anti-IL-6 receptor antibody treatment in ankylosing spondylitis. We consider that McH/lpr-RA1 is an adequate animal model of spondyloarthritis, and a promising tool for the evaluation and development of new treatment reagents and drug repositioning treatment in spondyloarthritis.

The results of IVIS analyses showed that there was a significant difference between the two groups in week 14 only. These results are inconsistent with the results of histological analyses, PCR and ELISA of SAA, which indicated significant differences after week 14. We consider that spontaneous arthritis in McH/lpr-RA1 mice becomes self-limiting around week 18. The histological findings showed significant differences in late phase arthritis, including bone erosion, pannus formation and ankylosis.

However, IVIS analyses did not indicate significant differences in late phase arthritis due to self-limitation of inflammation in the joints.

## Conclusion

In the present study, we have demonstrated for the first time that IL-6 signal blockade with MR16-1 significantly reduces the development of synovitis and joint destruction in the murine experimental model of spondyloarthritis, McH/lpr-RA1. Our results indicate that the progression of deformity associated with ankylosis continues even with anti-IL-6 receptor antibody treatment; indicating that further factors might be involved in the progression of joint ankylosis. McH/lpr-RA1 is a promising animal model for the elucidation of the mechanism of spondyloarthropathy and development of new treatment.

## Abbreviations

BV: Bone volume
CT: Computed tomography
cDNA: complementary DNA
ELISA: Enzyme-linked immunosorbent assay
ICG: Indocyanine green
IL: Interleukin
IVIS: In vivo bioluminescence imaging system
PBS: Phosphate-buffered saline
RT-PCR: Realtime polymerase chain reaction
SAA: Serum amyloid A
TNF: Tumor necrosis factor
TV: Total volume

## Declarations

## Acknowledgements

This study was supported by JSPS KAKENHI (18K09052) URL of Japan Society for the Promotion of Science is as follows: https://www.jsps.go.jp/english./index.html. The funders had no role in study design, data collection and analysis, decision to publish, or preparation of the manuscript.

## Author’s Contributions

Conceived and designed the experiments: TI, YM, SM, TK.

Performed the experiments: TI, YM, KT.

Contributed materials/analysis tools: YM, SM, TK Wrote the manuscript: TI, YM, NM, EI

## Ethics approval

Experiments were approved by The Tohoku University Animal Studies Ethics Committee.

